# A compound that directly and selectively stalls PCSK9 translation

**DOI:** 10.1101/083097

**Authors:** Nathanael G. Lintner, Kim F. McClure, Donna Petersen, Allyn T. Londregan, David W. Piotrowski, Liuqing Wei, Jun Xiao, Michael Bolt, Paula M. Loria, Bruce Maguire, Kieran F. Geoghegan, Austin Huang, Tim Rolph, Spiros Liras, Jennifer A. Doudna, Robert G. Dullea, Jamie H.D. Cate

**Author notes:** Current address: Pfizer Medicinal Chemistry, Cardiovascular, Metabolic and Endocrine Disease Research Unit, Pfizer Worldwide Research and Development, Cambridge, Massachusetts, 02139, USA. Correspondence should be addressed to R.G.D or J.H.D.C.

## Abstract

Proprotein Convertase Subtilisin/Kexin Type 9 (PCSK9) plays a key role in regulating the levels of plasma low density lipoprotein cholesterol (LDL-C). Here we demonstrate that the compound PF-06446846 inhibits translation of PCSK9 by inducing the ribosome to stall around codon 34, mediated by the sequence of the nascent chain within the exit tunnel. We further show that PF-06446846 reduces plasma PCSK9 and total cholesterol levels in rats following oral dosing. Using ribosome profiling, we demonstrate that PF-06446846 is highly selective for the inhibition of PCSK9 translation. The mechanism of action employed by PF-06446846 reveals a previously unexpected tunability of the human ribosome, which allows small molecules to specifically block translation of individual transcripts.

**One Sentence Summary:** A small-molecule PCSK9 inhibitor targets the human ribosome and selectively prevents PCSK9 synthesis.

## Introduction

Reduction of plasma low density lipoprotein cholesterol (LDL-C) through the use of agents such as statins represents the therapeutic standard of care for the prevention of cardiovascular disease (CVD),(1, 2) the leading cause of death in Western nations. Proprotein Convertase Subtilisin Kexin Type 9 (PCSK9) regulates plasma LDL-C levels by preventing the recycling of the LDL-receptor (LDLR) to the plasma membrane of hepatocytes.(3, 4) Humans with natural PCSK9 loss-of-function mutations display dramatically reduced LDL-C levels and decreased risk of CVD, yet display no adverse health effects(5-8). The robust LDL-C lowering observed with recently approved PCSK9 monoclonal antibodies (mAbs) when administered as a monotherapy or in combination with established LDL-C lowering drugs validates the therapeutic potential of inhibiting PCSK9 function(9-11). However, these therapeutic candidates require a parenteral route of administration, rather than being orally bioavailable. Utilizing a phenotypic screen for the discovery of small molecules that inhibit the secretion of PCSK9 into conditioned media, we have recently identified a compound family that inhibits the translation of PCSK9(*12*). However, the mechanism of translation inhibition exerted by these compounds remains unknown. Herein we describe a more optimized small molecule, PF-06446846 that demonstrates *in vivo* activity. We show that PF-06446846 induces the 80S ribosome to stall while translating PCSK9. We further demonstrate using ribosome profiling that despite acting through protein translation, a core cellular process, PF-06446846 is exceptionally specific, affecting very few proteins. The PF-06446846 mechanism of action reveals a previously unexpected potential to therapeutically modulate the human ribosome with small molecules as a means to target previously “undruggable” proteins.

## PF-06446846 inhibits PCSK9 translation by causing the ribosome to stall during elongation

The previously identified hit compound was adequate for initial *in vitro* characterization but *in vivo* assessment required improvements in pharmacokinetic properties(*12*). The potency, physicochemical properties and the off-target pharmacology associated with the hit compound were improved by structural changes to two regions of the molecule. These efforts led to the identification of compound PF-06446846 (Fig. 1A) with properties suitable for both *in vitro* and *in vivo* evaluation (Fig. S1 and Table S1). The synthesis and physiochemical characterization of PF-06446846 are described in the supplemental materials and methods, Figs. S2-S8 and Tables S2-S7. PF-06446846 inhibited the secretion of PCSK9 by Huh7 cells with an IC_50_ of 0.3 µM (Fig S1A). However, metabolic labeling of Huh7 cells with ^35^S-Met/Cys showed that decreases in PCSK9 were not a consequence of global inhibition of protein synthesis (Fig. S1B-C). Furthermore, proteomic analysis of the Huh7 cells utilizing stable isotope labeling with amino acids in culture (SILAC) indicated no general effect of PF-06446846 on the secreted and intracellular proteome (Fig S9, Supplementary Data Tables 1–3). Taken together, these results indicate that PF-06446846 exhibits a high degree of specificity for inhibiting the expression of PCSK9.

**Fig. 1.**
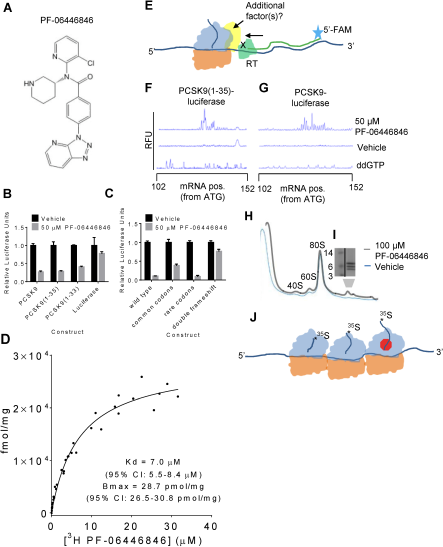
PF-06446846 targets the human ribosome inducing stalling during PCSK9 translation. **(A)** Structure of PF-06446846. **(B)** Luciferase activity of HeLa-based cell-free translation reactions programmed with mRNAs encoding PCSK9-luciferase, PCSK9(l-35)-luciferase, and PCSK9(1-33)-luciferase fusions and luciferase alone, in the absence (black bars) or presence (grey bars) of 50 |iM PF-06446846. **(C)** PF-06446846-sensitivity dependence on the amino acid sequence of PCSK9(1-33). PCSK9-luciferase fusions encode the native PCSK9 amino acid sequence with common codons or rare codons, or a native double-frameshifted mRNA sequence that results in a changed amino acid sequence (See Fig. S1E for sequences). All error bars represent one standard deviation of three replicates. **(D)** ^3^H-PF-06446846 binding to purified human ribosomes, K_d_: 7.0 μM (95% Cl: 5.5-8.4) B_max_: 28.7 pmol/mg (95% Cl: 26.5-30.8). The symbols within the graph represent the individual measurements obtained from three independent experiments. B_max_ and Kd values were calculated using GraphPad PRISM where the complete (n=) dataset was fit to the one site-specific binding equation. (E) Schematic of ribosomal toeprinting assays. 5’ 6-FAM labelled primers are extended by reverse transcriptase which terminates when blocked by a ribosome. **(F-G),** Electrophoreograms of toeprints of stalled ribosomes on the **(F)** PCSK9(l-35)-luciferase fusion construct and **(G)** full-length PCSK9-luciferase fusion. **(H)** Sucrose-density gradient profiles of cell-free translation reactions programmed with an mRNA encoding an N-terminally extended PCSK9, in the presence of 100 |iM PF-06446846 (grey) and vehicle (blue). (I) Tris-Tricine SDS-PAGE gels showing ^35^S-Met-labelled peptides that sediment in the polysome region of the gradient. **(J)** Model of the species isolated by density gradient centrifugation containing one stalled ribosome and two queued ribosomes.

To identify the specific mechanism responsible for translation inhibition by PF-06446846, we tested mRNAs encoding PCSK9-luciferase fusions in HeLa cell derived *in vitro* translation assays *(12).* PF-06446846 inhibited translation of PCSK9-luciferase fusion constructs containing only the first 35 residues of PCSK9, and displayed comparable activity towards the first 33 residues (Fig. 1B). In the HeLa cell-free translation assay, PF-06446846 inhibited PCSK9(1-35)-luciferase expression with an IC_50_ of 2 µM, while at the maximum concentration evaluated, 250 µM, the translation of luciferase without the PCSK9 N-terminal sequence was only inhibited by 20% (Fig. S1D). Translation of the protein fusion constructs was driven by the EMCV-IRES, indicating that PF-06446846 is unlikely to target PCSK9 translation initiation directly. When all the codons of PCSK9(1-33) were mutated to either common or rare synonymous codons (Fig. S1E), PF-06446846 still inhibited translation of PCSK9 (1-33)-luciferase (Fig. 1C), ruling out a role of the mRNA sequence. Conversely, PF-06446846 did not inhibit translation of a PCSK9(1-33) construct with two compensatory frameshifts that result in a near endogenous mRNA sequence but a non-endogenous amino acid sequence (Figs. 1C & S1E). These data indicate that PF-06446846 sensitivity is primarily dependent on the amino acid sequence of PCSK9. To further define the sequence requirements, we tested the activity of PF-06446846 against sets of N-terminal deletions, C-terminal deletions and alanine scanning mutations of PCSK9 (1-33). The most important regions in PCSK9(*1-33*) that confer sensitivity to PF-06446846 are Leu15-Leu20, residues 9-11 which include two tryptophan amino acids, and residues 31-33 (Fig. S10A-B). However, most mutations partially reduced the activity of PF-06446846, suggesting that multiple amino acid features of PCSK9(*1-35*) make contributions to its sensitivity to PF-06446846.

The sensitivity of the extreme N-terminal sequence of PCSK9 to PF-06446846 suggests that PCSK9(*1-35*) may act as a small-molecule-induced arrest peptide in the ribosome exit tunnel, similar to arginine attenuator peptide, TnaC, ErmCL and CatA86 (*13*). [^3^H]PF-06446846 binds to purified human ribosomes (K_d_= 7 µM) in filter-binding assays (Fig. 1D). In ribosomal toeprinting assays(14) of cell-free translation reactions programmed with full-length PCSK9 and PCSK9(*1-35*)-luciferase fusions, 50 µM PF-06446846 induced reverse-transcriptase early termination products consistent with stalling on and after codon 35 (Fig. 1F-G). The reverse transcription termination peaks were spread over 8-12 nucleotide (nt) positions, which could be due to a mixed population of ribosome complexes in which additional factor(s) are bound and block mRNA regions from reverse transcriptase access (Fig. 1E, yellow), or due to the compound causing the ribosome to stall at different codons. In cell-free translation reactions programmed with mRNA encoding full-length PCSK9 fused to a N-terminal extension, PF-06446846 also resulted in the appearance of polysomes containing three small radiolabeled peptides, suggesting that these PCSK9 nascent chains associated with one stalled ribosome, followed by two queued ribosomes (Fig. 1H-J). Most arrest peptides function only in one domain of life, i.e. solely in bacteria or solely in eukaryotes(*13*). In agreement with this phenomenon, PF-06446846 inhibited PCSK9(*1-35*)-luciferase translation in cell-free translation systems derived from rabbit reticulocytes, wheat germ and yeast but not from *E. coli* (Fig. S10C-F).

## PF-06446846 reduces circulating PCSK9 and total plasma cholesterol levels *in vivo.*

To explore the safety of the compound and to gain insight into the *in vivo* activity of PF-06446846, male rats were orally administered PF-06446846 at doses of 5, 15 and 50 mg/kg daily for 14 days. Plasma PF-06446846 (Table S8) and PCSK9 concentrations were measured at 1, 3, 6 and 24 hours following the first and twelfth dose and total plasma cholesterol levels were assessed in fasted animals just prior to necropsy on Day 15. Dose dependent lowering of plasma PCSK9 was observed following single and repeated dosing of PF-06446846 (Fig. 2A-B). In addition to the reduction in circulating levels of PCSK9, evidence of inhibition of PCSK9 downstream function was observed at the 50 mg/kg dose, with a statistically significant 30% decrease in total plasma cholesterol and 58% decrease in LDL cholesterol but no significant decrease in high-density lipoprotein (HDL). This encouraging effect on circulating PCSK9 concentration was achieved in the absence of modulating plasma liver function markers, with no treatment related changes observed for alanine transaminase (ALT), aspartate transaminase (AST) or albumin (Figs. 2C-E S11). Although reductions in cholesterol were observed, this endpoint is poorly assessed in rodents and the clinical experience shows that reduced free PCSK9 levels represent the primary determinant of the improved lipid profile in human *(5-9, **11**)*.

**Fig. 2.**
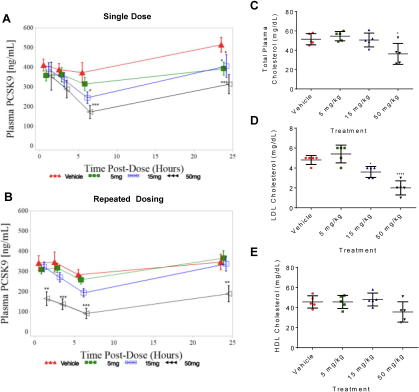
Oral administration of PF-06446846 reduces plasma PCSK9 and total cholesterol levels in rats. **(A-B)** Plasma PCSK9 levels following **(A)** a single and **(B)** 12 daily, oral doses of PF-06446848. Rats were administered the indicated dose of PF-06446846 and plasma concentrations of PCSK9 were measured by commercial ELISA at 1, 3, 6 and 24 hours after dosing (A) or the twelfth daily dose (B). Symbols represent mean concentration +/- standard error and were jittered to provide a clearer graphical representation. Data were analyzed using a mixed model repeated measure (MMRM) with Treatment, Day, Hour as fixed factors, Treatment by Day and Hour as an interaction term, and Animal as a random factor. Significant level was set at a level of 5%. No adjustment for multiple comparisons was used. *p<0.05, **p<0.01, ***p<0 **ooi. (C-E)** Total plasma **(C),** LDL **(D),** and HDL **(E)** cholesterol levels in rats measured 24-hours following 14 daily oral doses of PF-06446846. Symbols represent individual animal values. The middle horizontal bar represents the group mean +/- standard deviation. Difference between group means relative to vehicle was performed by a 1-way ANOVA followed by a Dunnett's multiple comparisons test; * p<0.05, **** p<0.0001.

During the two-week dosing period, PF-06446846 was tolerated. There was a small decrease (11-13% relative to vehicle) in food consumption at 50 mg/kg PF-06446846 that was not associated with any changes in body weight. Histological and clinical chemistry examination of samples collected at day 15 indicated no dose-limiting changes compared to vehicle at the 5 and 15 mg/kg dose level. Administration of PF-06446846 at 50 mg/kg/day demonstrated a minimal decrease in bone marrow cellularity (primarily involving erythroid parameters), that correlated with a decrease in red cell mass (9%). Furthermore, mild reductions in white blood cells (52%), neutrophils (40%) and lymphocytes (54%), and reductions in the T cell (approximately 54% in the total, help and cytotoxic T cells) and B cell populations (58.4%), as well as minimal necrosis of the crypt cells of the ileum (in 1 of 5 animals), were observed at 50 mg/kg/day. Importantly, no histopathological findings were observed at any dose of PF-06446846 in the liver, the organ responsible for the majority of PCSK9 production as well as maintaining whole-body cholesterol homeostasis(15, 16).

## Translation focused genome-wide identification of PF-06446846 sensitive genes

While the absence of adverse histopathological changes is consistent with the high degree of selectivity observed in SILAC experiments (Fig. S9), we used ribosome profiling(17-19) to provide a higher resolution understanding of the selectivity of PF-0644846 at the level of mRNA translation. Huh7 cells were treated with 1.5 and 0.3 µM PF-06446846 (5X and 1X the IC_50_ in Huh7 cells) (Fig. 1A) or a vehicle control, for 10 and 60 minutes in three biological replicates, and ribo-seq libraries were prepared. To measure mRNA abundance and translational efficiencies we subsequently conducted a second ribosome profiling study in which Huh7 cells were treated with 1.5 µM PF-06446846 or vehicle for one hour and both ribo-seq and mRNA-seq libraries were prepared from the same sample.

Metagene analysis(20, 21) of the distribution of ribosomal footprints relative to the start and stop codons displayed the hallmarks of ribosome footprints(18, 19), including 3-nt periodicity through the coding DNA sequence (CDS) regions and a minimal number of reads mapping to 3’-UTR regions (Figs. 3A-D and S12). We also observed high correlation between replicates in reads aligning to individual genes (Fig. S13). We consistently observed a compound-independent depletion of reads aligning to the first forty codons of the CDS regions, a large buildup of reads on the stop codon, and a small queued ribosome peak upstream from the stop codon(22) (Fig. 3A-D and S12). This buildup may be due to omission of the cycloheximide pre-treatment step*(22-24),* which we omitted to avoid an artefactual buildup of reads near the start codon(19). PF-06446846 treatment induced no change in the average distribution of reads along the CDS after 1 hour of treatment (Fig. 3C) and only a small PF-06446846-induced accumulation of reads mapping to the first 50 codons after a 10 minute treatment (Fig. 3A), indicating that PF-06446846 does not cause pausing or stalling on most transcripts. The PF-06446846-induced increase in early read counts at the 10 minute timepoint, but not at the 60 minute timepoint, may indicate the induction of cellular stress upon the initial exposure to PF-06446846 followed by adaptation within the one hour timeframe.

**Fig. 3.**
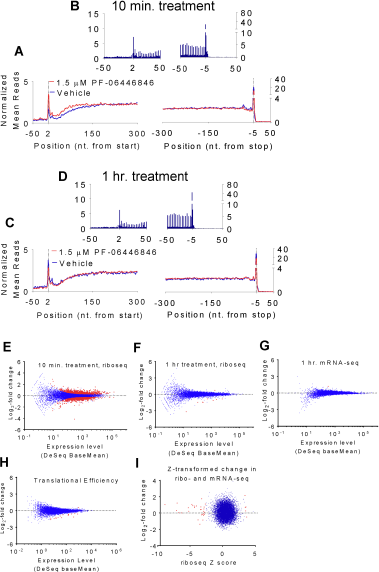
PF-06446846 does not cause widespread ribosomal stalling. **(A-D)** Metagene analysis of ribosomal footprint distribution relative to the start and stop codons for cells treated with 1.5 |iM PF-06446846 (red trace) and vehicle (blue trace) for **(A,B)** ten minute treatment and **(C,D)** one hour treatment. **(A,C)** Values are averaged over three nucleotides for clarity. **(B,D)** Zoom to show three nucleotide periodicity and specificity for CDS regions. The normalized mean reads (NMR) were calculated as in ref (20) The normalized read count at a given position on a particular mRNA is the number of reads aligning to that position divided by the average read density along the CDS. These values are then averaged across all transcripts of sufficient length. **(E-G)** Log2-fold change in read counts plotted against expression levels for the **(E)** 10-minute treatments, **(F)** 1 hour treatments **(G)** mRNA-seq datasets for the TE data. Genes with a significant (FDR < 10%) change in expression are highlighted in red. **(H)** TE plotted as in e-g. Red points indicate Z-score >3.0 when compared to genes of similar expression levels(22) (See materials and methods for Z-score transformation).

Differential expression analysis using the ribosome footprint data revealed 1956 genes are differentially expressed after 10 minutes of treatment at a false discovery rate (FDR) of 10% (DeSeq) but only 9 after one hour (Fig. 3E-F). Only a single gene, EFNA3, displayed a difference in transcript levels (FDR < 10%). mRNA-seq and translational efficiency (TE) analysis reveals that almost all of the change in expression levels occurs at the translation step (Fig. 3G-I). The two features present in the data at 10 minutes but not at one hour, the relative global depletion of reads near the 5’ end of the CDS regions and the large number of genes displaying a brief change in translation levels, could be indicative of cellular stress after ten minutes of treatment. Thus, we focused the following analyses generally on the 1-hour treatment time.

Examination of ribosomal footprints aligning to the PCSK9 transcript (Fig. 4A-C) revealed a PF-06446846-dependent buildup of footprints centered on codon 34, consistent with ribosomal stalling at this position. The number and distribution of fragments from mRNA-seq was unaffected by PF-06446846 (Fig. 4D). PF-06446846 treatment also resulted in a decrease in ribosome footprints mapping 3’ to the stall site on the PCSK9 transcript. Notably, the magnitude of the decrease in footprints 3’ to the stall site is similar to the level of inhibition of PCSK9 expression as measured by ELISA (Fig. 4E), indicating that for PF-06446846-sensitive proteins, the decrease in reads mapping 3’ to the stall site could function as a surrogate measurement of the extent of translation inhibition for a given protein. Thus, PF-06446846 targets should be detectable through differential expression analysis using only the reads that align 3’ to the stall site. In our second ribosome profiling study, we found that the magnitude of the buildup of reads at the stall site was smaller for both PCSK9 (Fig 4C) and other targets, while the number of reads mapping 3’ to the stall site remained consistent. The variability in magnitude of the stall peak could be an artefact of the size selection step during library preparation; subsequent to our initial set of experiments, it was reported(25, 26) that ribosome profiling of stalled ribosomes can yield a broader range of protected footprint sizes than the conventional 26-34 nt range(*19*).

**Fig. 4.**
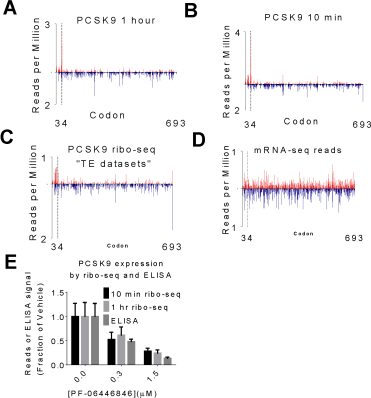
The PF-06446846-induced stall site is revealed by ribosome profiling. **(A-D)** Ribosome footprint density plots for the PCSK9 coding region from Huh7 cells treated for **(A)** 1 hour **(B)** 10 minutes. **(C)** ribo-seq datasets from the second study, **(D)** mRNA-seq datasets from the second study. The upward red bars indicate readmaps from cells treated with 1.5 μM PF-06446846 and the blue downward bars represent vehicle. **(E)** The footprint density downstream from the stall in the 10 minute treatment (black) and hour minute treatment (light grey) compared with PCSK9 expression as measured by ELISA (middle grey). Error bars represent one standard deviation of three replicates.

To identify and quantify the sensitivity of individual proteins to PF-06446846, we adopted a computational approach to identify transcripts that could potentially have PF-06446846-induced stalls, estimate the position of the stalls, and use differential expression analysis using only reads aligning 3’ to the stall site (Fig. 5A). To estimate a 3’ bound for the positions of potential PF-06446846-induced pauses, we adapted the approach previously reported (21). For each gene, we plotted the percentage of the total reads aligning at or 5’ to each codon to generate cumulative fractional read (CFR) plots (Fig. 5B-C). If PF-06446846 induced a stall, the CFR plot should increase rapidly 5’ to the stall and level off 3’ to the stall. We define the maximum divergence between the PF-06446846 and vehicle CFR plots as the D_max_ and the codon at which D_max_ occurs as the D_max_ position, which is analogous to the KS position previously described (21). For genes with one or more PF-06446846-induced stalls the D_max_ position occurs 3’ to the last stall site. For genes with a high D_max_ (Z-score > 2) we used reads mapping 3’ to the position of D_max_ for differential expression analysis (Fig. 5D, see methods for details). Using this approach, we identified 22 PF-06446846-sensitive genes at the 60 minute timepoint (Figs. 5 E-I and S14, Table S9) and 44 genes at the 10-minute timepoint (Dmax Z-score > 2, DeSeq FDR > 10%). With the exception of CDH1 and IFI30, all PF-06446846-sensitive proteins identified at the one hour timepoint were also identified at the 10-minute timepoint. To test the robustness of our approach we also analyzed the data from the second ribosome profiling experiment with the same pipeline. Despite most stall peaks being smaller in the second study, we identified all of the same PF-06446846-sensitive sequences as for the data from study 1, demonstrating the advantage of using information from the entire transcript instead of a single codon.

**Fig. 5.**
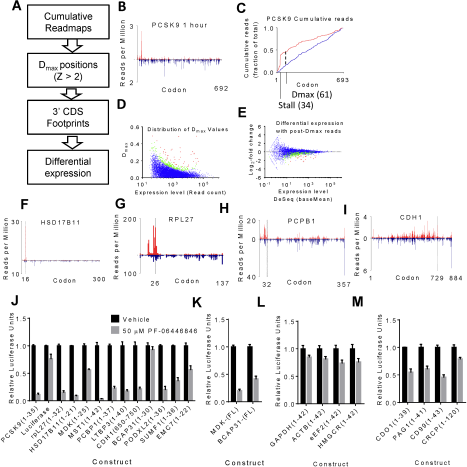
Identification and validation of PF-06446846-sensitive nascent chains. (A) Outline of the approach to identify PF-06446846 targeted mRNAs. (B) Example readplot and (C) Example cumulative fractional read (CFR) plot for PCSK9. A CFR plot depicts at each codon the percentage of reads aligning at or 5ʹ to that codon. In all plots data from 1.5 μM PF-06446846 treatments are shown in red and vehicle treatments are shown in blue. The major stall site is denoted with *, the position of D_max_ is marked. **(D)** Scatterplot showing the distribution of D_max_ values as a function of read counts, red indicates D_max_Z-score > 3 (see materials and methods for Z-score calculations) and green indicates 2 < Z-score < 3. (E) Scatterplot of fold change vs expression level when reads mapping 3’ to D_max_ position (for Z-score > 2) or codon 50 (for Z-score < 2) are used. Genes for which D_max_ Z-score > 2 and DeSeq FDR < 10% are highlighted in red, for Dmax Z-score > 2 but FDR > 10% in green, for D_max_ Z-score < 2 with FDR < 10% in purple. **(F-I),** Example readplots for PF-06446846-sensitive protiens, **(F)** HSD17B11, **(G)** RPL27, **(H)** PCBP1, and **(I)** CDH1. Bars representing the treatment dataset are red and go upwards and bars representing the vehicle datasets are blue and go downwards. All graphs are derived from the 1-hour treatment time in the first study. **(J)** Cell-free translation assays showing inhibition of translation by 50 μM PF-06446846 when the stall sites identified by ribosome profiling are fused to the N-terminus of luciferase. **(K)** Inhibition of *in vitro* translation of full-length Midikine— and BCAP31-luciferase fusions in the cell-free translation system. **(L)** *In vitro* translation of control constructs not predicted to be inhibited by PF-06446846 from cell-based experiments. **(M)** *In vitro* translation of constructs with PF-06446846-induced stalls identified only at the 10-minute treatment time.

SILAC-based proteomic analyses of lysates of Huh7 cells treated for 4 h or 16 h with 0. 25 |μM or 1.25 |μM PF-06446846 (the same cells as those from which the secretome data were obtained) failed to detect most of the hits identified by ribosome profiling including PCSK9, when data were analyzed using the same criteria as for the secretome (Fig. S9E-H). With a reduction in stringency (proteins with 3 unique peptides accepted), a 2- to 3-fold reduction in cadherin-1 (CDH1) production was detectable after 16 h treatment with PF-06446846 (Fig. S9G-H). These results suggest that depletion of the levels of transcript-stalled proteins are likely to occur at a rate governed by protein turnover.

In addition to stalling on the main open reading frame (ORF), it is possible that stalling on upstream ORFs could lead to a change in translation of the main ORF. We thus analyzed all upstream ORFs for a change in read density and found none that displayed an increase in reads with PF-06446846 treatment that was consistent across replicates. SORCS2, FAM13B and LRP8, the three genes that are downregulated at the 1 hour timepoint but do not have a Dmax Z-score > 2 do not have translated uORFs.

To validate our approach, we tested the translational inhibitory activity of PF-06446846 for a set of the targets identified in the 1-hour datasets in HeLa-derived cell free translation reactions. In the all but two cases, translation of constructs consisting of the predicted stall site fused to luciferase was inhibited by PF-06446846 (Fig. 5J). For the other two proteins, Midikine and BCAP31, the translation of luciferase fusions to the full-length proteins was inhibited by PF-06446846 (Fig. 5K). Four control sequences predicted not to be PF-06446846-sensitive were inhibited at comparable levels as luciferase alone (Fig. 5L).

We next tested the translation inhibition of PF-06446846 towards four example “stall sequences” identified only in the 10-minute dataset. PF-06446846 inhibited translation of all of these transcripts only slightly more than luciferase alone (Fig. 5M). These results indicate that the most sensitive PF-06446846 targets are those identified using the 1-hour treatment time. The additional effects seen in the 10-minute treatment could be due to a partial adaptation of the cells on the 1-hour timescale, or the cells could be sensitized during the first 10 minutes of compound treatment due to the sudden media changes required for the experiment. In either case, these data indicate that 1-hour treatment times are most appropriate for evaluation of this and similar compounds.

As a complimentary approach, we also calculated the change in mean read position or center of density *(22)* between the 1.5 µM PF-06446846 and vehicle data, and used a Z-score transformation *(27)* to identify outliers. 17 transcripts with a significant change in the center of read density (Z > 3) were identified in the one hour datasets (Figs. 6A and S14T). Of these, 13 were also identified using the D_max_ approach (Fig. S14T). For the hits identified using only center-of-density analysis, VPS25 and TM2D3 had stalls but no decrease in downstream reads, indicating that the stall was unlikely to result in a decrease in protein production (Fig. S14P-Q). MAPRE1 did not have a clear stall site (Fig. S14R). COX10 (Fig. S14S) had a series of PF-06446846-induced stalls near the stop codon which would be difficult to detect using the D_max_ approach because of a lack of downstream reads to quantify. We also used center of density analysis to confirm the bias for stalling near the 5’ ends of the CDS by repeating the center of density analyses omitting the first 50, 100 and 150 codons respectively (Fig. 6B-D). In all cases, the cluster of outliers disappeared, indicating that, while there are exceptions, PF-06446846-induced stalls most often occur in the first 50 codons.

**Fig. 6.**
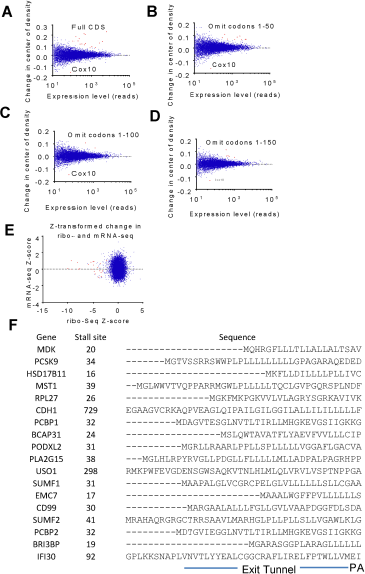
Features of PF-06446846-sensitive transcripts. **(A)** Changes in the mean read position or center of density(22) plotted against expression levels. Genes with a local Z-score greater than 3 are highlighted in red. **(B-D)** Center of density analysis with first **(B)** 50, **(C)** 100 and **(D)** 150 codons omitted, showing that stalling preferentially occurs before codon 50. (E) Expression changes for PF-06446846-sensitive transcripts occur at the step of translation. Plot of Z-score transformed (22) read counts for mRNA-seq and ribo-seq. PF-06446846-sensitive nascent chains are highlighted in red. **(F)** Alignment of PF-06446846-sensitive sequences. The sequences are aligned according to the stall position and the residues predicted to reside in the ribosome exit tunnel, P-site and A-site are indicated.

To confirm that the decrease in reads downstream from PF-06446846 induced stalls occur as a result in changes in translation as opposed to mRNA levels, we plotted the change Z-score transformed(22) (see materials and methods) in mRNA-seq reads versus the changes in ribo-seq reads (Fig. 6E). For all identified PF-06446846-sensitive sequences, the changes in ribosome footprints, were due solely to changes at the level of translation (Fig. 6E). We additionally tested how transcript levels of PF-06446846 targets change with longer-scale treatments and under different growth conditions by treating Huh7 cells grown in either standard or lipoprotein-depleted FBS for four and 24-hours with 1.5 µM PF-06446846, then measuring the transcript levels by RT-qPCR. With four biological replicates, no consistent changes in target transcript levels were found (Fig. S15A-B), consistent with PF-06446846-induced changes in protein expression occurring at the translation step.

## Discussion

To our knowledge PF-06446846 is the first example of an orally-active small molecule that inhibits PCSK9 directly. Although the narrow safety margin likely prevents clinical development of PF-06446846, it demonstrates that it may be possible to modulate PCSK9 function with a small molecule. This has been a challenge for small molecules by conventional means as it requires the disruption of the small flat interaction between PCSK9 and the LDL-receptor (LDLR) that represents the primary contact patch at neutral pH. The importance of PCSK9 in regulating LDL-C clearance by LDLR is underscored by the greater degree of LDL-C lowering possible with the mAbs relative to statins. However, by their nature, the anti-PCSK9 mAbs are unable to modulate PCSK9 and LDL-C levels in an intermediate fashion. Inhibition of PCSK9 synthesis by a small molecule could provide additional options for patients and physicians by offering a range of inhibition to balance safety and efficacy.

This work also represents the first demonstration of the potential for selective inhibitors of eukaryotic mRNA translation as a therapeutic approach. The reduction of PCSK9 demonstrated by PF-06446846 *in vivo* with no sign of toxicity in the liver is consistent with the high selectivity observed by ribosome profiling. It is unclear whether a lack of complete selectivity evident in the very small number of transcripts whose translation was perturbed by PF-06446846 is responsible for the adverse effects seen outside the liver at the highest dose.

PF-06446846 induces the ribosome to stall on the PCSK9 transcript a few codons beyond to the end of the signal peptide. Ribosome profiling reveals that despite acting on the human ribosome, PF-06446846 is exceptionally specific. Interestingly, the validated off-targets have few common features in their primary structures predicted to be present in the ribosome exit tunnel when stalling occurs (Fig. 6F).Although a number of the PF-06446846 sensitive sequences include a leucine repeat or hydrophobic stretch N-terminal to their stall sites, the vast majority of hydrophobic stretches in all of the translated proteins are unaffected. These results indicate that, although PF-06446846 sensitivity is determined by amino acid sequence, stalling cannot be predicted by a simple primary structure motif. Additional requirements for stalling are likely to include the structure adopted by the nascent chain in the exit tunnel(28, 29) or the context in which the sensitive sequence occurs.

One common feature of the PF-06446846-sensitive proteins is that stalling usually occurs early in the protein coding region; 15 of the 18 stalls identified in the one-hour treatment occur before codon 50, with the most N-terminal stall site occurring at codon 16. The heightened sensitivity of N-terminal regions of proteins to PF-06446846 could be due to a unique feature early in the elongation phase of translation. Most stalls would likely occur before the nascent chain emerges from the ribosomal exit tunnel or when only a few residues are extruded. Our mutagenic analysis also indicates that the most critical residues for PF-06446846-induced stalling reside in the exit tunnel, pointing to the ribosomal exit tunnel as a critical element of PF-06446846 action. This is consistent with previous findings that the ribosomal exit tunnel acts as a functional environment, allowing certain peptides and small molecules to induce ribosome stalling(13, 30-35).

Most translation regulation mechanisms in eukaryotes identified to date act on the initiation step(36). PF-06446846 presents a case of protein production rates controlled during elongation. Translation regulation during elongation presents an interesting mechanistic puzzle as, theoretically, every ribosome that initiates should eventually result in a full-length protein, regardless of how long it takes to traverse the CDS. Our mRNA-seq and RT-qPCR data rule out the possibility that the stalled ribosome induces No-Go decay (37) or other mRNA quality control mechanisms that decrease overall mRNA-levels. It is possible that PF-06446846 induced stalling near the N-terminus could impede initiation, thus lowering the overall initiation rate. Evidence for “queued” ribosomes (Fig. 1H-J) suggest impeding of initiation could occur for stalls near the N-terminus. However, an impact on translation initiation could not explain the effects of more C-terminal stalling on CDH1 levels (Fig. S9G-H), and would predict that PF-06446846-sensitive transcripts would display a bias for higher TE, which they do not (Fig S15C). Alternatively, the stalled ribosomes may be removed from the transcript by a “rescue” mechanism and not complete translation. Future experiments will be required to determine the underlying mechanisms of protein reduction due to ribosome stalling by PF-06446846.

While PF-06446846 directly and selectively inhibits the translation of PCSK9 during the elongation phase, most drugs and drug candidates known to modulate human translation target translation initiation factor complexes or upstream signaling pathways such as mTOR(38, 39). Furthermore, they generally modulate translation of a large number of mRNAs. One drug candidate, Ataluren, selectively induces read-through of premature stop codons while not affecting termination at native stop codons(40). The present example of PF-06446846 demonstrates that drug-like small molecules have the potential to directly modulate human translation elongation for therapeutic purposes. The variability in the protein primary structures affected by PF-06446846 suggest that each target may be affected by a slightly different mechanism and that analogues of PF-06446846 could be further optimized for improved selectivity for PCSK9 or even to target other proteins.

## Acknowledgments

We thank Jiang Wu for contributions to the SILAC study as well as Beijing Tan and Christopher Holliman for their bioanalytical support. We thank Santos Carvajal-Gonzalez for statistical analysis support. This work used the Vincent J. Coats Genomics Sequencing laboratory at the University of California, Berkeley, supported by National Institutes of Health (NIH) S10 Instrumentation Grants S10RR029668 and S10RR027303, as well as computing resources of the Center for RNA Systems Biology (NIH grant P50-GM102706). The authors thank Brian Samas for the X-ray structure determination of PF-06446846 and Chris Limberakis and Steve Coffey for synthetic intermediates. The authors wish to thank Pfizer Medicinal Chemistry and the Pfizer Emerging Science Fund for their support.

## Author Contributions

A.L., D.P., L.W., J.X. contributed to the chemical synthesis. N.G.L., B.M. and D.P. designed and performed the biochemical experiments. N.G.L. prepared the ribosome footprint libraries. D.P and K.G. performed and analyzed the SILAC experiments. N.G.L. and A.H. analyzed the high throughput sequencing data. N.G.L. K.F.M., P.L., M.B., T.R., S.L., J.A.D., R.G.D and J.H.D.C. designed the study, analyzed the data and wrote the manuscript.

## Competing financial interests

N.G.L., D.P., A.L., D.W.P., L.W., J.X., M.B., P.M.L., B.M., K.F.G., A.H., K.F.G., A.H., K.F.M., R.G.D. and S.L. are employees of Pfizer, Inc.

## Supplementary Materials

Materials and Methods

Figures S1-S15

Tables S1-S9

Supplementary Data Tables 1-4

Additional Files S1-S16

